# Red blood cell lingering modulates hematocrit distribution in the microcirculation

**DOI:** 10.1101/2022.08.16.504126

**Authors:** Yazdan Rashidi, Greta Simionato, Qi Zhou, Thomas John, Alexander Kihm, Mohammed Bendaoud, Timm Krüger, Miguel O. Bernabeu, Lars Kaestner, Matthias W. Laschke, Michael D. Menger, Christian Wagner, Alexis Darras

## Abstract

The distribution of red blood cells (RBCs) in the microcirculation determines the oxygen delivery and solute transport to tissues. This process relies on the partitioning of RBCs at successive bifurcations throughout the microvascular network and it is known since the last century that RBCs partition disproportionately to the fractional blood flow rate, therefore leading to heterogeneity of the hematocrit (i.e. volume fraction of RBCs in blood) in microvessels. Usually, downstream of a microvascular bifurcation, the vessel branch with a higher fraction of blood flow receives an even higher fraction of RBC flux. However, both temporal and time-average deviations from this phase-separation law have been observed in recent works. Here, we quantify how the microscopic behavior of RBCs lingering (i.e. RBCs temporarily residing near the bifurcation apex with diminished velocity) influences their partitioning, through combined *in vivo* experiments and in silico simulations. We developed an approach to quantify the cell lingering at highly-confined capillary-level bifurcations and demonstrate that it correlates with deviations of the phase-separation process from established empirical predictions by Pries *et al.* Furthermore, we shed light on how the bifurcation geometry and cell membrane rigidity can affect the lingering behavior of RBCs, e.g. rigid cells tend to linger less than softer ones. Taken together, RBC lingering is an important mechanism that should be considered when studying how abnormal RBC rigidity in diseases such as malaria and sickle-cell disease could hinder the microcirculatory blood flow or how the vascular networks are altered under pathological conditions (e.g. thrombosis, aneurysm).

## Introduction

Partitioning of red blood cells (RBCs) through bifurcations of blood vessels determines how the oxygen is delivered to tissues and organs. Early works from Zweifach-Fung and Fåhræus showed that it is far from being a trivial phenomenon [1, 2, 3], and since then it has been an area of intensive research [4, 5, 6, 7, 8, 9, 10]. More accurately, vessels with a higher flow rate tend to collect an even higher proportion of RBCs. This fact is known as the Zweifach-Fung effect and leads to inhomogeneity in the hematocrit of various vessels [1]. An empirical model which reproduces this average behavior has been developed by Pries *et al.* [11, 12, 13, 14]. However, recent works, either experimental or computational, have shown that deviations from this model can arise for various reasons [7, 8, 15]. In particular, it has been shown that RBC partitioning at bifurcations in the smallest vessels of the microvascular network tend to deviate from this empirical model prediction [11, 16, 17, 18, 19]. On the other hand, recent numerical simulations and *in vivo* experiments highlighted that RBCs tend to linger at the apex of bifurcations, modifying the dynamics of cells entering a single vessel and their characteristic inter-distance [20, 17, 21]. In the present study, we systematically analyzed how the lingering of RBCs influence their partitioning in the microcirculation, through complementary *in vivo* experiments and *in silico* simulations. We show that deviations from the Pries model are directly correlated with normalized average duration of cell lingering. This asymmetric drive can either reinforce or revert the Zweifach-Fung effect. Moreover, RBCs exhibit large deformations while lingering. We further explored numerically how rigidified cells differ from healthy ones in their lingering and partitioning. Our results demonstrate and quantify the lingering as a source of deviation from the Zweifach-Fung partitioning, and provides a new explanation of how pathologically rigidified cells can impair the partitioning of RBCs in the microcirculation [22, 23, 24, 25, 26, 27, 28].

## Materials and Methods

### In vivo experiments

#### Permissions

Experiments were performed in Syrian golden hamsters according to the German legislation on protection of animals and were approved by the local governmental animal protection committee (permission number: 25/2018). Hamsters were maintained on a standard 12/12 h day/night cycle and water and food were provided ad libitum. Detailed methods are described below for each type of investigation.

#### Animal preparation and microscopy

Hamsters with an age of 5 to 7 weeks, weighing 55 to 70 g, were used for the implantation of a dorsal skinfold chamber [29]. The surgery was performed under deep anesthesia (150 mg/kg ketamin (Serumwerke Bernburg AG)/ 0.25 mg/kg domitor (Orion Pharma) i.p.), with intraoperative pain medication by carprofen (10 mg/kg, Zoetis, s.c.). A previously described protocol [29] was followed.

Briefly, the back of the hamsters was shaved and a titanium chamber consisting of two frames was implanted on the lifted dorsal skinfold. On the frame with a circular observation window (diameter of 10 mm) the cutis, subcutis and retractor muscle were removed to expose the striated skin muscle for later observation of the microcirculation. The window was closed with a cover glass that was fixed with a snap ring. An implanted dorsal skinfold chamber is depicted in Fig. 1(*a*). Animals were allowed to recover for 72 hours after the procedure.

**Figure 1:**
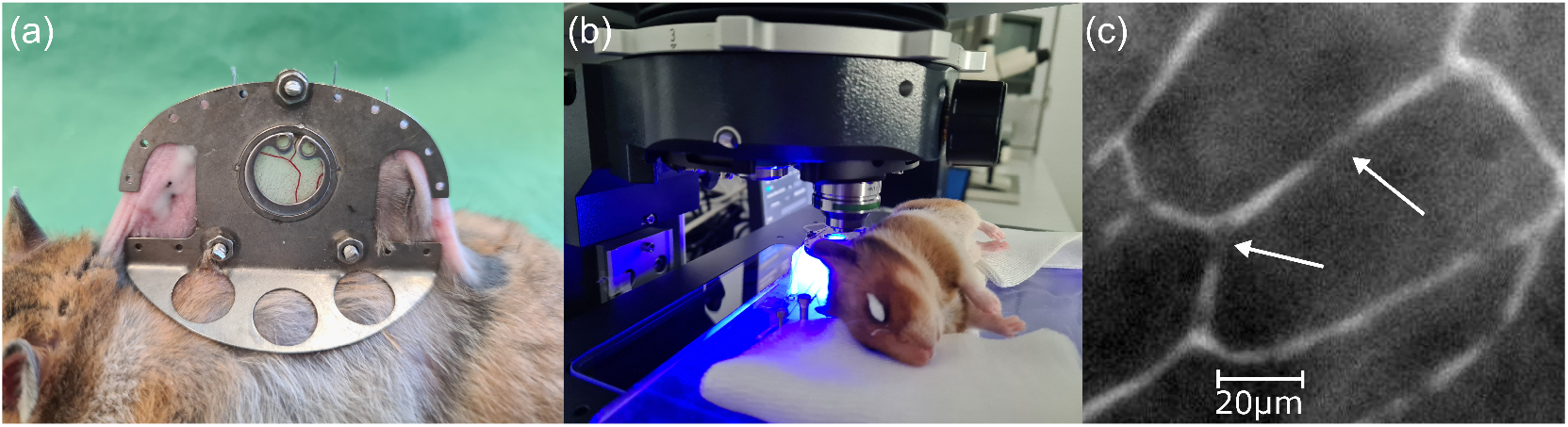
Experimental Methods: (*a*) Dorsal skinfold chamber implanted on the back of a hamster. (*b*) Anesthetized hamster placed underneath the objective of an epifluorescence microscope. (*c*) Example of a microvascular network imaged by fluorescence microscopy. The dyed plasma appears bright, while RBCs in the capillaries and the surrounding tissues are dark (arrows).

Hamsters were anesthetized as described above prior to intravital microscopy. A volume of 100 *μ*L of the fluorescent plasma marker fluorescein isothiocyanate(FITC)-labeled (5%, 150 kDa, SigmaAldrich) was injected retro-orbitally and the animals were fixed on a plexiglas stage, as illustrated in Fig. 1(b). Several capillary bifurcations in different areas of the chamber window were chosen for epifluorescence microscopy (Axio Examiner A1, Zeiss). FITC-labeled dextran was excited with a 480 nm LED (Colibri 7, Zeiss) and image contrast was additionally enhanced by simultaneously using transmitted blue light that is absorbed by hemoglobin, making RBCs appear darker. Imaging was performed with 20x (LD A-Plan, NA=0.35, Zeiss), 50x (LD EC Epiplan-Neofluar, NA=0.55, Zeiss) or 100x (LD C Epiplan-Neofluar 100x, NA=0.75, Zeiss) long-distance objectives. Video acquisition was carried out with a digital camera (Hamamatsu Orca Flash 4.0, C13440) using the software ZEN 3.1 Blue (Zeiss). For each video, 2000 frames were recorded at different rates according to flow velocity (100 to 170 Hz). A 2×2 binning was applied during acquisition. An example of an obtained image is given in Fig. 1(*c*).

#### Image analysis and lingering quantification

Despite the animal being fixed on the plexiglas stage, its breathing and/or muscles movement caused slight translations of the microscopic field of view in the image sequence (Supp. Fig. S1(*a*)). In order to determine the motion of the RBCs in the vessels, we first correct those translations by translating each image by the maximum of its 2D-correlation with the first image in the sequence. This process creates a movie where the vessels and surrounding tissues are still (see Supp. Fig.S1(b)). A Gaussian filter (with standard deviation of 2 pixels) is then applied to despeckel the images, and a mask is drawn around the vessels. The creation of this mask is facilitated by also considering the image showing the standard deviation of each pixel intensity along time, which is higher in the vessels with transient (dark) RBCs and offers a guide to the eye for the mask drawing (see Supp. Fig.S1(c)). The diameters *D* of the various vessels are computed as twice the average distance from the skeleton pixels of the masks to the border of the vessels, using standard Matlab functions bwmorph and bwdist (see Supp. Fig.S1(c)). The image series is then inverted and binarized inside the mask area with a threshold based on a user-defined fraction of the Otsu threshold [30]. Otsu thresholding method was used because the average intensity of images is usually slightly flickering. However, the automated threshold algorithm overestimated the threshold required to detect the whole RBCs, hence we implemented a user-defined ratio to adjust it (see Supp. Fig.S1(d)).

A tracking algorithm was then used to determine the velocity of the detected RBCs [31]. The variations of light intensity along the vessels, in conjunction with the collisions and transient aggregations of RBCs, didn’t allow us to reliably follow all RBCs along their entire trajectories. However, as we focused on bifurcations where the mother vessel (*M*) and the two daughter vessels have a hematocrit low enough to distinct single RBCs, this tracking allowed us to extract the spatial distribution of the RBC velocity (i.e. we performed Particle Tracking Velocimetry), on which we performed our further analyses. In this manuscript, we will refer to both daughter vessels as main and secondary daughter vessels, respectively abbreviated *MD* and *SD*. The *MD* is identified as the daughter vessel with higher fractional blood flow rate.

More accurately, in order to detect lingering of RBCs, we applied a method similar to previous investigations [21]: We defined a circle with a diameter of 6 *μ*m (i.e. one cell’s main diameter) around the bifurcation apex (Fig. 3(*a*)). The cells were considered to linger if their center of mass was in this circle and had a velocity lower than the local minimum detected in the PDF of the velocity (Fig. 3(*b*)). The velocity PDF displayed in Fig. 3(*b*) was obtained when considering the mother vessel and the bifurcation area.

### Numerical simulations

Complementary simulations for the RBC flow in representative capillary bifurcations were run with *HemeLB* (https://github.com/hemelb-codes/hemelb) using the immersed-boundary-lattice-Boltzmann method following our previous approach [16]. First, three-dimensional (3D) luminal surface models were reconstructed from binary masks of the capillary bifurcations using open-source software *Pol-Net* [32], assuming circular cross-sections of varying diameter along each vessel. Then the flow domain enclosed by the luminal surface was uniformly discretized into cubic lattices of fine voxel size Δ*x* = 0.25 *μm*, aimed at resolving cell dynamics with high resolution at the bifurcation apex. To initialize the simulation, inflow/outflow boundary conditions based on experiment-measured time-average volume flow rates were imposed at the end of the mother vessel (*M*) and the main daughter branch (*MD*, the daughter branch with higher flow rate) of each bifurcation, and a reference pressure for the secondary daughter branch (*SD*, the daughter branch with lower flow rate). No-slip boundary conditions were imposed at the vessel wall. The simulation time step length was Δ*t* = 1.04 × 10^*−*6^*s*.

The simulation for each bifurcation was initialized with (RBC-free) plasma flow. Once the plasma flow became converged, RBCs were randomly inserted from the *M* vessel through a cylindrical flow inlet with constant feeding hematocrit (*H*_F_) as measured from experiments. When RBCs reached the end of *MD* or *SD*, they were removed from the simulation domain. Each RBC was modeled as a cytosol-filled capsule with an isotropic and hyperelastic membrane consisting of 5120 triangular facets. The mechanical properties of the RBCs were governed by elastic moduli governing different energy contributions, such as shearing and bending of the membrane. The cytosol was treated as a Newtonian fluid with the same viscosity as the suspending medium, *i.e.* blood plasma. The viscosity of the RBC membrane itself was not considered. Two different RBC sizes (*D*_rbc_ = 6 *μm* and 8 *μm*) and membrane rigidity levels (stain modulus *κ_s_* = 5 × 10^*−*6^, 5 × 10^*−*5^*N/m* for representing a normal and a hardened cell) were considered.

## Results and Discussion

### Determination of flow rates and expected Zweifach-Fung partitioning

In order to determine how the lingering of RBCs influence their partitioning, we used the empirical model of Pries *et al.* for the Zweifach-Fung effect [13] as a baseline. For this purpose, we first determined the blood flow rate *Q* in each vessel. Since the vessels were smaller than the main radius of the RBCs, we assumed a plug flow. We also assumed that the difference between the tube and discharge hematocrits was negligible in our observations. The flow rates within all bifurcation segments were therefore initially assessed from the velocities (V) and diameter (D) measurements of the individual segments

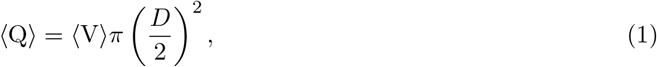

similarly to previous works [11, 33, 21]. Mass conservation stipulates that the total volume flow rate from the mother vessel *Q_M_* is divided into the daughter branches and satisfy *Q_M_* = *Q_MD_* + *Q_SD_*. The subscripts *MD* and *SD* refer to the Main Daughter and Secondary Daughter, respectively. To obey this conservation law and minimize experimental errors, values obtained from Eq.(1) were corrected following the normalization process described by Pries *et al.* in previous similar measurements [11]. Similarly, the measured erythrocyte flow rates *E_i_* = *H_i_Q_i_*, *H_i_* being the local hematocrit, were also corrected to obey *E_M_* = *E_MD_* + *E_SD_*. The empirical law for the Zweifach-Fung effect developed by Pries *et al.*[11, 12, 13] describes the relation between fractional blood flow, *FQ_b_*, and the fractional erythrocyte flux, *FQ_e_*, in the microvascular bifurcation. The fractional blood flows *FQ_b_* for both daughter vessels were calculated by dividing the flow rate *Q* in the respective daughter by the flow rate in the mother vessel. The fractional erythrocyte flow rates *FQ_e_* were computed as the ratio of the corrected erythrocytes flow rates, e.g. *E_MD_/E_M_* for the main daughter. With these definitions, the model from Pries *et al.*[11, 12, 13] states

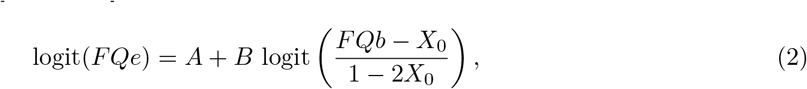

where 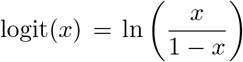 and *A*, *B*, *X*_0_ are related to the geometry of the bifurcation and the hematocrit in the mother vessel

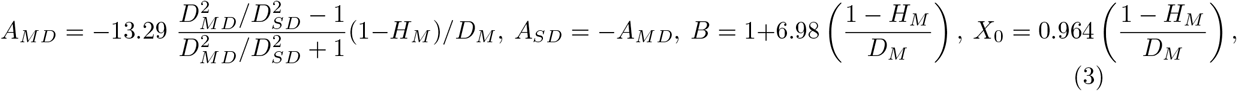

where *H* refers to the measured hematocrit, assessed as the average area fraction of RBC in the mask during the experiment [12, 13]. The subscripts indicate in which vessel the quantity is considered.

### Lingering and partitioning

#### Global description

##### Experiments

Figure 2(*a, b, c*) shows characteristic bifurcations (two Y-shaped and one T-shaped) where all required parameters could be extracted from the image sequences. We calculated the empirical predictions by Eq.(2) for each available bifurcation, and defined the deviations from the model as *δZF* = |Δ*FQ_e_*(*EX*) − Δ*FQ_e_*(*ZF*)|. We noted Δ*FQ_e_* = *FQ_e_*_(*MD*)_ − *FQ_e_*_(*SD*)_ the difference of fractional erythrocyte flow rate, and the expressions *EX* and *ZF* refer respectively to the experimental result and the empirical prediction. The fractional blood flow rate *FQ_b_* is given by the experimental measurements, and the predicted fractional erythrocyte flow rate *FQ_e_*(*ZF*) is computed through Eq.(2) (see Fig. 4(*a*) for a schematic).

**Figure 2:**
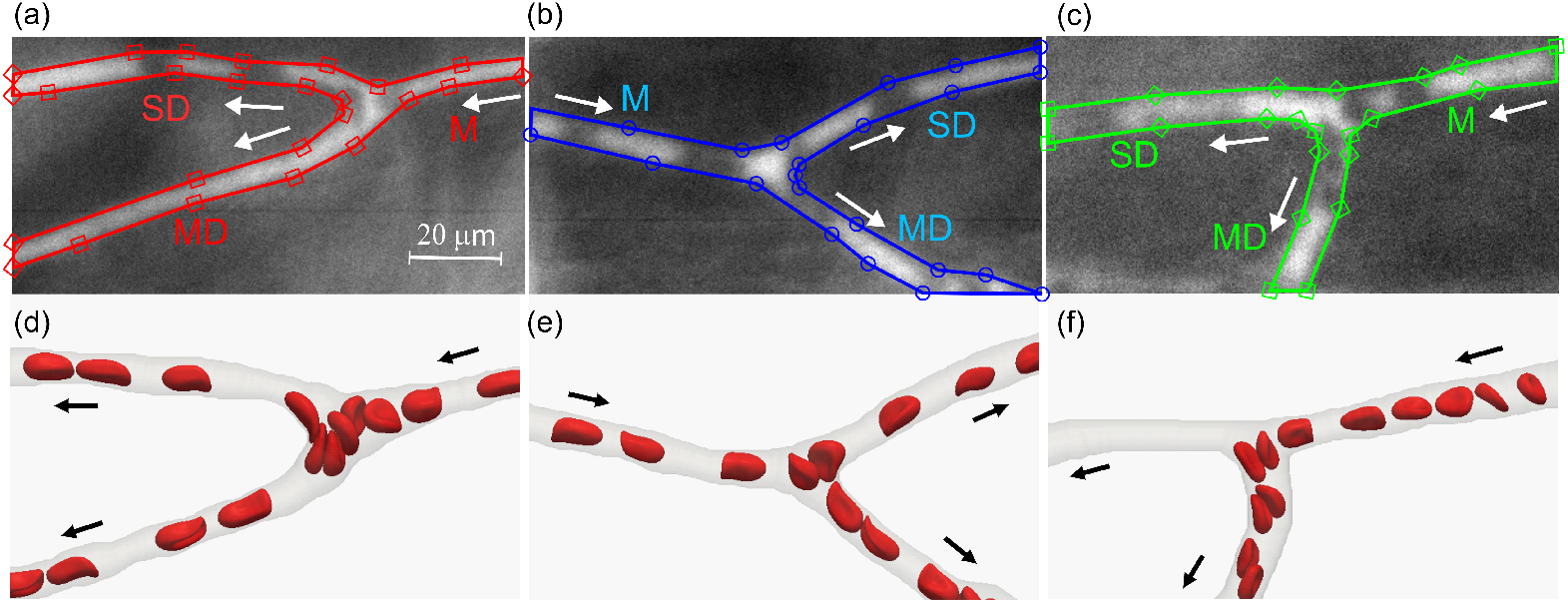
Representative bifurcations in experiments and simulations. (*a, b, c*) Three bifurcations selected from the experiments hereafter referred to as “BIF-a”, “BIF-b”, “BIF-c”, respectively. The mother vessel (M), the main daughter (MD) and secondary daughter (SD) are annotated and the arrows indicates the flow direction. The plasma was stained with fluorescent dye, which is bright in the images. Dark areas in the vessels indicate the RBCs. The border of the masks used to analyze the bifurcations are depicted in colored symbols. Consistent colors and symbols are used for these bifurcations throughout the article. (*d, e, f*) Simulated RBC flow in reconstructed bifurcations (resp. (*a, b, c*)), as characterized experimentally.

In order to quantify the lingering of RBCs in each bifurcation, we first determined the average duration of the lingering events ⟨*τ*⟩. The total lingering time was considered as the sum of the duration of all detected lingering events. We then determined the ratio of this total lingering time to the number of the RBCs as the characteristic average lingering time ⟨*τ*⟩. Since this time is highly correlated with the flow rate and the transit time of a RBC through the bifurcation (i.e. the red circle defined in Fig. 3(*a*)), we normalized it by *t_s_* = *R/*⟨*V_M_* ⟩. The radius *R* = 3 um is the radius of the red circle defined in Fig. 3(*a*) and *V_M_* the average velocity in the mother vessel. The time *t_s_* is then a characteristic advection time that a cell would need to go through the lingering detection area. We defined the ratio of these two characteristic times as *Pe_λ_* = ⟨*τ*⟩/*t_s_*. The ratio *Pe_λ_* defines the equivalent of a Péclet number, since it compares two characteristic times of different transport modes in the bifurcation (lingering and advection). As shown in Fig. 4(*c*), the deviation *δZF* from the Zweifach-Fung prediction Eq.(2) strongly correlates with *Pe_λ_* (Pearson correlation coefficient *r* = 0.811, with a *p*-value of *p* = 0.007).

**Figure 3:**
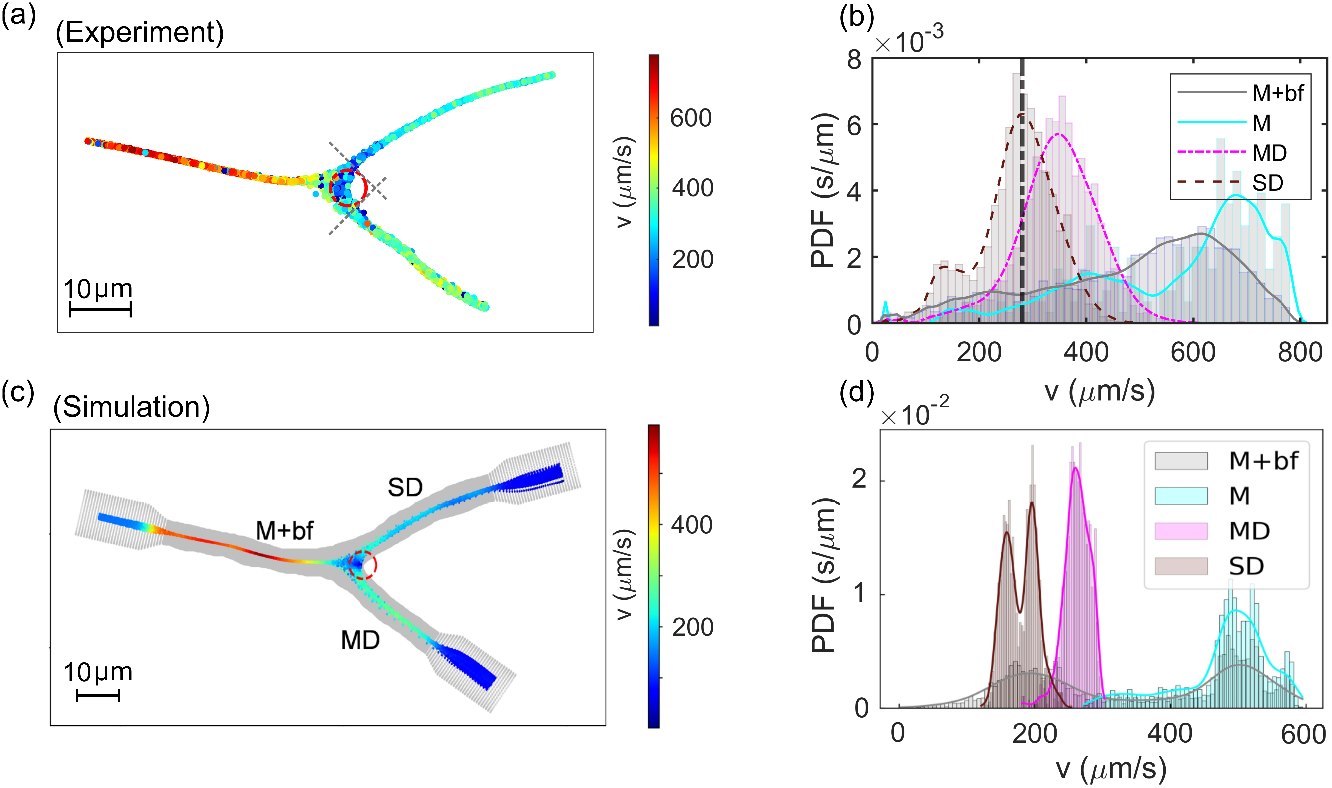
RBC velocity distributions in experiments and simulations. (*a*) Spatial distribution of RBC velocities obtained through particle tracking in experiment for “BIF-b”. Lingering is considered to occur if a cell is inside the red circle (6 μm, size of a typical mouse RBC) with a velocity lower than the threshold velocity, which is determined as a local minimum in the probability density function (PDF) of velocities obtained from the mother vessel and the bifurcation area (M+bf), as shown in panel (*b*). For comparison, the velocity PDFs for the main and secondary daughter branches (MD and SD), and the mother vessel (M) are also shown in (b). (*c*) Corresponding color map of RBC velocities measured along simulated cell trajectories in “BIF-b”. (*d*) Velocity PDFs measured in different vessel branches of the simulation as for the experimental data in (b).

**Figure 4:**
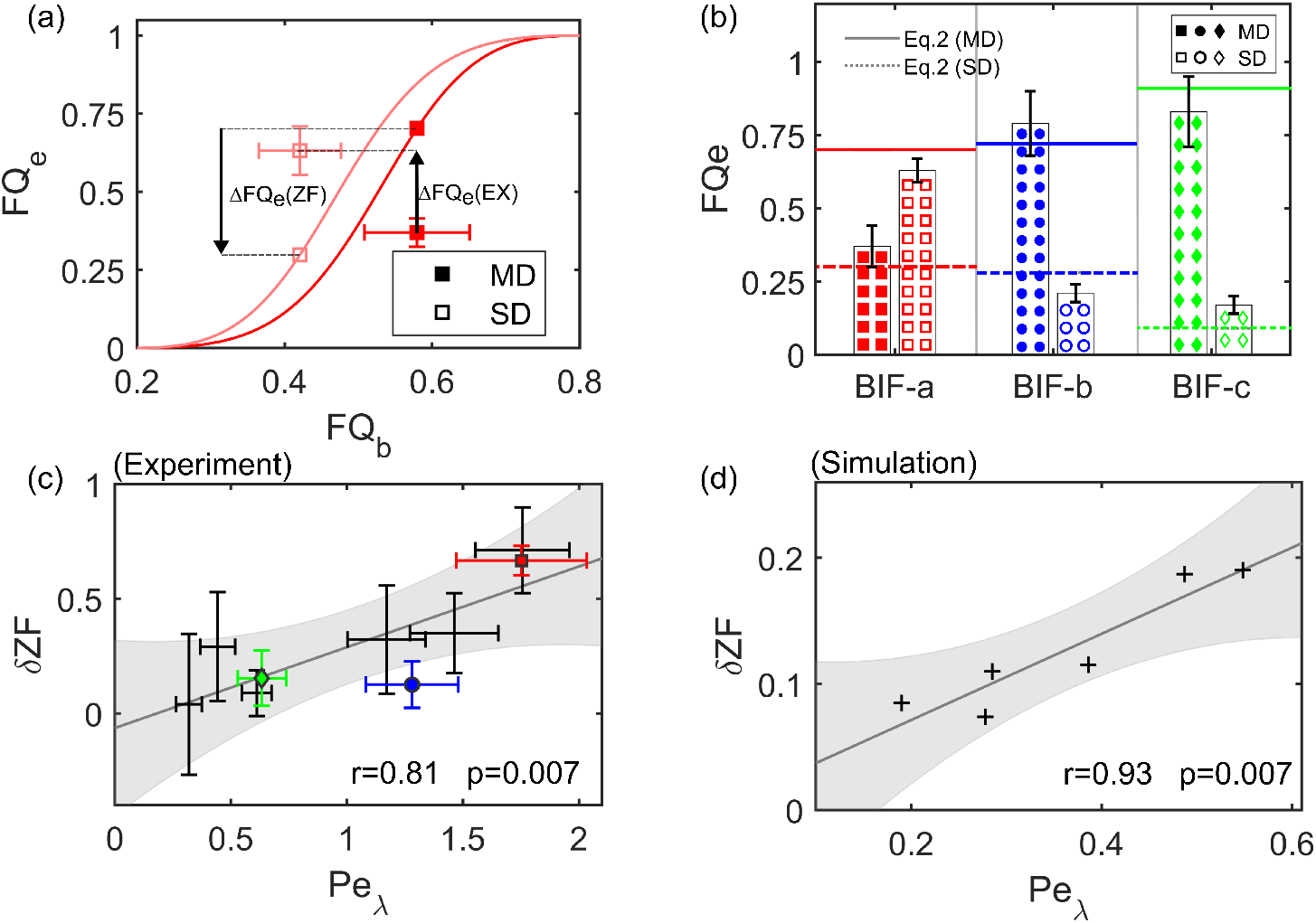
Experimental and numerical results on RBC partitioning versus lingering. (*a*) Comparison of experimental RBC fluxes against the empirical predictions (for “BIF-a” as in Fig.2(*a*)). The axes *FQ_e_* and *FQ_b_* are for the fractional RBC flux and fractional blood flow, respectively. We calculated deviation from the Zweifach-Fung prediction as *δZF* = |Δ*FQ_e_*(*EX*)− Δ*FQ_e_*(*ZF*)|. (*b*) The bars show experimental results against the lines, which are empirical predictions by Eq.(2). The data here are for the three characteristic bifurcations in Fig. 2(*a, b, c*), with consistent colors and symbols. Dashed lines and hollow symbols refer to the secondary daughter vessel (with lower fractional flow rate *FQ_b_*). Note the inversion of the Zweifach-Fung effect for “BIF-a”. (*c*) Experimental deviation *δZF* from Eq.(2) as a function of the lingering Péclet number *Pe_λ_*. The points are experimental data, and the solid line shows linear regression fitting of the data points, with the shaded area indicating 95% confidence interval prediction from the fit. All data gathered during this study (including, but not limited to, “BIF-a” “BIF-b” “BIF-c” in Fig. 2(*a, b, c*)) are included in this graph. The Pearsonr correlation coefficient is 0.81, with a *p*-value of 0.007. (*d*) Correlation between *δZF* and *Pe_λ_* for simulation data. The crosses are simulation data and the solid line shows linear regression fitting of the data points. The Pearson-r correlation coefficient is 0.93, with a *p*-value of 0.007.

Interestingly, we observed in some case that the deviation from the Zweifach-Fung effect caused by the lingering can revert the partitioning. Indeed, in the case of the (red) characteristic bifurcation highlighted in Figure 2(*a*), the main daughter (with higher *FQ_b_*) is also the branch with higher *FQ_e_*. This is often referred to in the literature as reversed partitioning [7, 8, 15]. In our data, we only observed such reverse partitioning for the highest *Pe_λ_* ≈ 1.75.

##### Simulations

The three characteristic bifurcations from experiments (see Fig. 2(*a c*)) were also simulated under equivalent flow conditions and RBC volume fractions. The phenomenon of RBC lingering was reproduced (Fig. 2(*d – f*)). To quantify the lingering behavior, equivalent procedure to the experimental data analysis was applied to analyze the simulation data.

In line with the experimental particle tracking velocimetry (PTV) measurements, the cell velocities diminished as the RBCs approached the bifurcation apex (Fig. 3(*c*)). After entering one of the daughter branches (*MD* or *SD*), the RBCs accelerated and their instantaneous velocity increased. These patterns were well captured by the velocity PDFs compiled for the individual vessel branches (Fig. 3(*d*)). Quantification of lingering events based on the RBC velocities revealed that the percentages of lingering cells for the three bifurcations were 48%, 84% and 32%, respectively. For hardened RBCs with a membrane strain modulus ten times as large, the ratios become 31%, 68% and 35%.

The correlation between the lingering intensity of RBCs and their partitioning at the bifurcations, namely *Pe_λ_* and *δZF*, were evaluated similarly as for the experimental data. A strong association was found, indicating a linear increase of *δZF* against *Pe_λ_* (Pearson-r correlation coefficient 0.93, *p* < 0.01). Nevertheless, the absolute magnitude of *δZF* and *Pe_λ_* were substantially smaller (but in a proportional manner) than their experimental counterpart in Fig. 4(*c*). This discrepancy may arise from simulation configurations that are different from *in vivo* experiments, *e.g.* the absence of endothelial surface layer (ESL, with a thickness of roughly 0.4-0.5 *μ*m) and RBC glycocalyx in the numerical model. In any case, the correlation between the lingering Péclet number *Pe_λ_* and the deviation from the Pries model was qualitatively similar in the experiment and the simulation.

#### Lingering asymmetry in experiments

For insights into how the lingering behavior could modify RBC partitioning at the bifurcation, we further examined its symmetry between the two daughter branches. Indeed, as can be seen from Supplementary Movie 1, the RBCs tend to enter the lower daughter branch (*MD*) almost only if there is another cell already lingering at the bifurcation apex. Due to cell collisions at the apex, it was not possible to precisely determine the proportion of lingering cells entering each daughter merely by analyzing their trajectories. Therefore, for each daughter separately, we evaluated which cells detected were lingering by monitoring RBCs in a thin mask extracted from the daughter (Fig. 5(*a*)). We then checked if there was a lingering event at the bifurcation (also monitored in a thin mask) at a time *t_a_* earlier for each cell detection, where *t_a_* accounts for the average advection time of the cells between the bifurcation and the mask in the daughter. This advection time was determined by calculating the convolution of the temporal hematocrit in the daughter mask and that in the bifurcation mask (see Supp. Fig.S2(*b*)).

**Figure 5:**
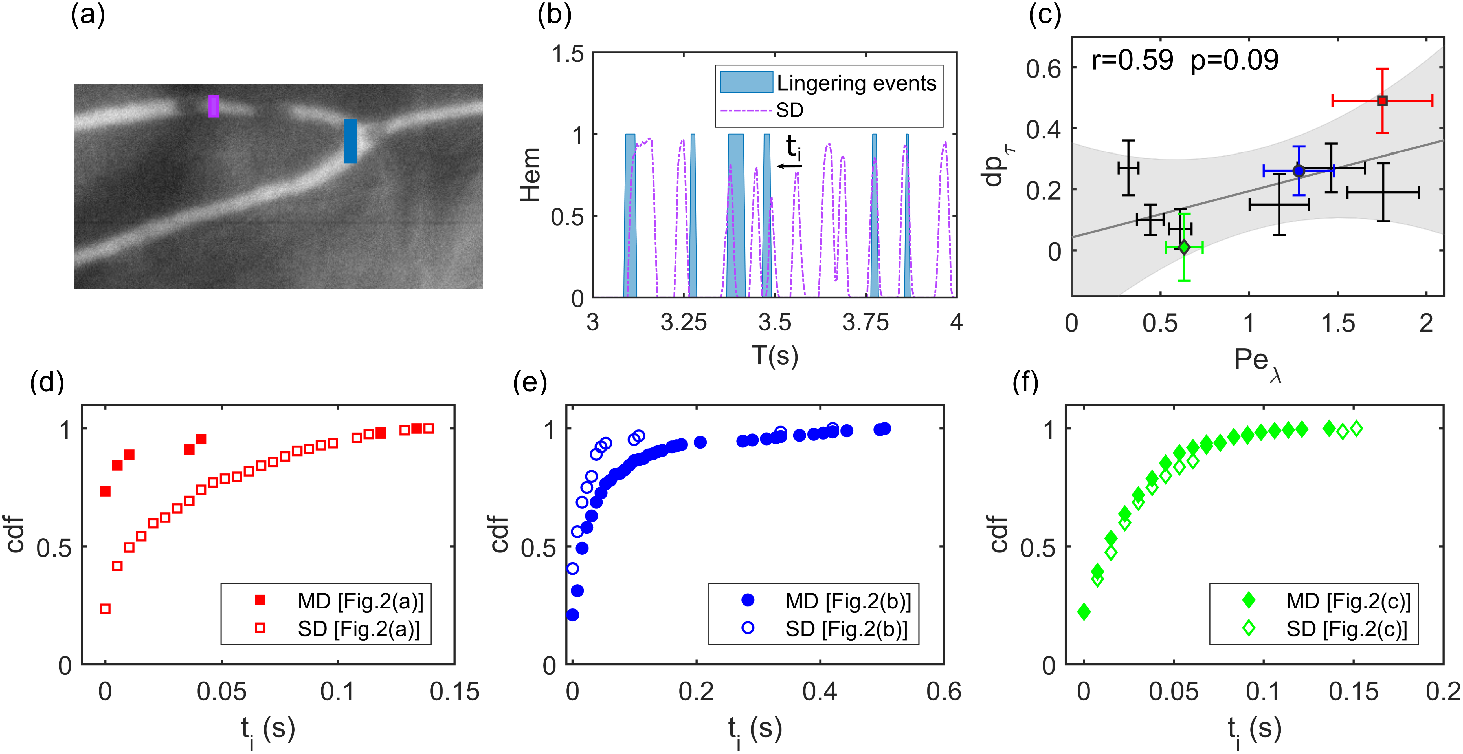
Asymmetry in RBC lingering at the bifurcation. (*a*) Example of mask set used to determine the proximity time *t_i_* of RBC transits against lingering events. The (blue) mask in the bifurcation is used to determine the average advection time of cells to the (magenta) mask in the secondary daughter (*SD* branch) via convolution of the hematocrit in both masks (see main text). Equivalent analysis is performed independently for the *MD* branch (not shown here). (*b*) Extraction of the proximity time *t_i_* of each cell. The hematocrit variation over time, extracted from the daughter’s mask, is first translated by the average advection time *t_a_* (see Supp. Fig.S2). Then, for each peak of this hematocrit along time, a proximity time *t_i_* is obtained as the temporal distance from the closest lingering event. This process is performed independently for the two daughters. The correlation between experimental *dP_τ_* (difference in the proportion of lingering cells in SD, MD calculated from CDFs at *t_i_* = 0) and *Pe_λ_*. The Pearson-r correlation coefficient is 0.59 (*p* = 0.09). (*d, e, f*) Cumulative Density Function (CDF) of the proximity time to lingering events for both daughters of the characteristic bifurcations. When a significant difference (considerable *dP_τ_*) is observed in the experiments, higher proportion of lingering cells (and a closer proximity with lingering events) is usually obtained in the daughter branch with the lower fractional RBC flux *FQ_e_*.

We realized that for such sorting, some lingering events at the bifurcation could not be attributed to any advected cell into the daughters. Indeed, the advection time of each cell can vary slightly, depending on its interactions with other cells near the apex. After translated by *t_a_*, the cell transits in the bifurcation can be temporally shifted towards the lingering event, but not exactly located within its bandwidth. For more accurate characterization, we therefore measured the proximity time *t_i_* between the closest lingering event and the transit time (detection time in the daughter minus *t_a_*) for each cell (Fig. 5(*b*)). Cells with a proximity time *t_i_* = 0 can then be determined as lingering. We further defined *dp_τ_* = |*p_SD_* − *p_MD_*| as the difference between the proportion of lingering cells in the two daughter branches (Fig. 5(c)). Because cells with a small proximity time *t_i_* might also have been lingering or might have interacted with another cell lingering at the apex, we used the difference between the first two points of the CDF distribution of *t_i_* (Fig. 5(*d*−*f*)) as errorbars for *p_SD_* and *p_MD_*. As can be seen in Fig. 5(*d*−*f*), the proximity time *t_i_* distribution and the proportion of cells identified as lingering are distinct for the two daughters under large *Pe_λ_*. For the experimentally observed cases as highlighted in Fig. 5(c), the asymmetry in the proportion of lingering cells is weakly correlated with the lingering intensity *Pe_λ_*. In simulation, both *dp_τ_* and *Pe_λ_* for the simulated cases (three in total) are smaller than their experimental counterpart, but in a proportional manner with qualitative agreement (see Supp. Fig.S2(*c*)).

### Origin of lingering in experiments

In order to understand what causes the cells to linger, we identified geometrical features of the bifurcations correlating with the lingering Péclet number *Pe_λ_*. Due to the definition of the lingering events as a significant reduction of the RBC velocity in the bifurcation, it is likely that cells linger at the stagnation point, i.e. the point where the local plasma velocity vanishes. At this position, the cells will also experience a negligible drag-force, therefore not pushing the cell towards any daughter vessel. The position of the stagnation point, which is also the intersection of the bifurcation’s wall and the flow divider plane, depends on the fractional flow rate of the daughter branches [34]. The distance of the stagnation point from the center line *L_s_*, which is depicted in the Fig. 6(a), can be approximated through

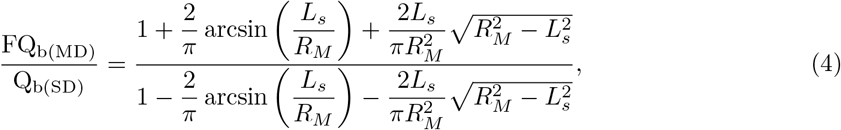

where *R_M_* is the radius of the mother branch (see Supplemental Material for further justification). For the situations where numerical simulations were compared to the experimental data, location of the stagnation point determined this way were located within one pixel of distance from the position indicated by the streamlines (See Supplemental Fig. S3). This approximation can then be considered to be consistent with the available numerical data.

**Figure 6:**
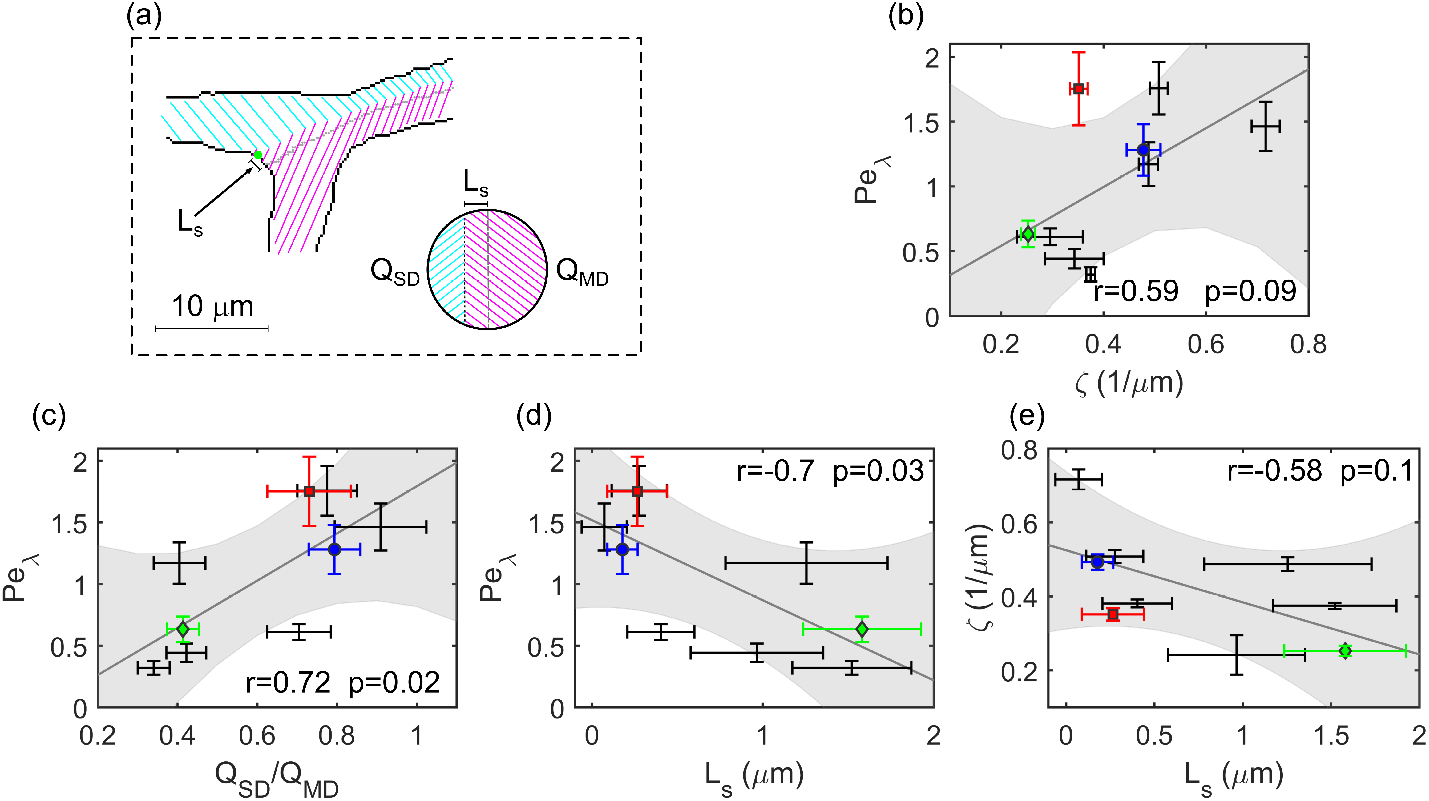
Potential origin of RBC lingering. (*a*) Schematic highlighting the flow split and the stagnation point at bifurcations. Flow in the mother branch splits into flows *Q_MD_* and *Q_SD_* in the MD and SD branches, respectively. The stagnation point is indicated by the green point, whose distance from the apex *L_s_* is highlighted with the arrow and scale symbol. The filled circle shows an idealized cross-section of the mother vessel with simplified and flat flow separatrix. (*b*) Correlation of the lingering Péclet number *Pe_λ_* with the curvature of the bifurcation at the stagnation point. (*c*) Correlation between *Pe_λ_* and the flow rate ratio between the secondary and main daughters *Q_SD_/Q_MD_*. (*d*) Correlation between *Pe_λ_* and the distance between the stagnation point and the bifurcation apex *L_S_*. (*e*) Correlation between *L_S_* and the curvature at the stagnation point *ζ*. In (*b − e*), the points are experimental data, the solid lines show linear regression fitting of the data points, and the shaded areas indicate the 95% confidence interval prediction from each fit.

Since the cells at the stagnation point experience only a negligible drag force, one can assume that the dominating forces on the cells arise from their interaction with the endothelial layer. In turn, this interaction probably depends on the geometry of the wall at this location, which can mostly be described through its curvature. In order to find the curvature of each bifurcation, we used the bestfit circle function at the stagnation point (Supp. Fig. S3 illustrates this process). We defined the curvature of the bifurcation as the inverse of the radius of the locally fitting circle and showed that it correlates with *Pe_λ_* (*r* = 0.59 and *p* = 0.09) (see Fig. 6(b)). Interestingly, when testing correlations with further possibly influencing parameters, we found that the lingering Péclet number *Pe_λ_* also correlates with the ratio of the flow rates between the secondary and main daughters *Q_SD_/Q_MD_* (Fig. 6(c)). Since this parameter determines the distance *L_S_* between the apex of the bifurcation and the stagnation point through Eq.(4), this parameter *L_S_* also correlates with *Pe_λ_*. Furthermore, the bifurcations where the stagnation point is further from the apex generally have a lower curvature at the stagnation point, as shown by Fig. 6(e).

We therefore hypothesize that the flow rate ratio determines where the cells can linger, while the curvature of the endothelial layer at this lingering position determines the intensity of the lingering.

The lingering Péclet number would then be determined by an interplay between global parameters of the bifurcation (*Q_SD_*/*Q_MD_*) and local geometries of the endothelial layer (the curvature at the stagnation point, *ζ*). When it comes to the experimental correlations, the fact that a lower p-value is found between *Pe_λ_* and *ζ* than *Pe_λ_* and *Q_SD_*/*Q_MD_* likely arises from the fact that the determination of *ζ* also relies on the approximation for *L_S_* and then the measurement of *Q_SD_*/*Q_MD_*. The error propagation on these successive quantities then likely decreased artificially the correlation between *Pe_λ_* and the successively determined parameters *L_S_* and *ζ*.

More geometrical parameters, which failed to demonstrate a significant correlation with *Pe_λ_*, are included in the Supplementary Materials (see Supp. Fig. S4).

### Effect of cell rigidity and hematocrit in simulations

Besides the curvature of the stagnation point at the apex, other factors can also affect the interaction between the cells and the bifurcation, leading to either weakened or enhanced RBC lingering. Taking a typical Y-type bifurcation for example (Fig. 7(*a*)), three physiologically-relevant aspects are explored: the cell rigidity *κ*_s_, the feeding Hematocrit *H*_F_ and the cell size *D*_RBC_ (*D*_rbc_ = 6 *μ*m unless otherwise specified).

**Figure 7:**
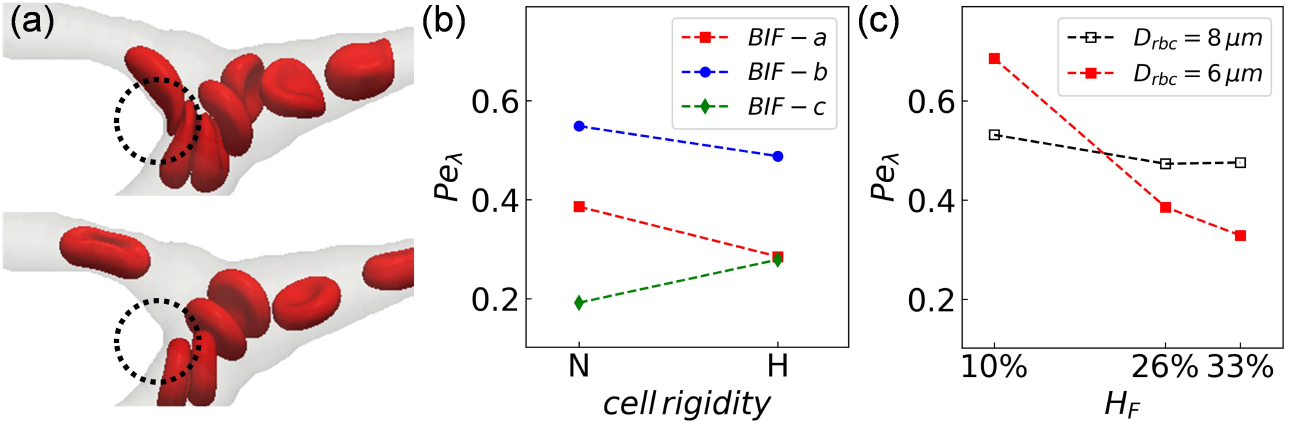
Effect of cell rigidity and feeding hematocrit on the lingering of RBCs. (*a*) Steric repulsion for (top) normal “N” and (bottom) hardened “H” RBCs at the apex of a Y-type bifurcation corresponding to “BIF-a” in Fig. 2. (*b*) *Pe_λ_* versus cell rigidity for normal and hardened RBCs at “BIF-a” ‘BIF-b” ‘BIF-c” (see Fig. 2), *D*_rbc_ = 6 *μ*m. (*c*) *Pe_λ_* versus feeding hematocrit *H*_F_ at “BIF-a” for normal RBCs of two different sizes *D*_rbc_ = 6, 8 *μ*m.

Increased cell rigidity (imitating diseased RBCs with higher-level stiffness) at the Y-type bifurcation is found to introduce strong cell-wall and cell-cell steric repulsion at the apex (see enlarged gaps near the branching point in Fig. 7(*a*)), thus reducing the overall lingering frequency and average duration of individual lingering events as indicated by a smaller lingering Péclet number *Pe_λ_* (“BIF-a”, Fig. 7(*b*)). The same is found for “BIF-b” too (Fig. 7(*b*)). Contrarily, for a T-type bifurcation (less common bifurcation type in microvascular networks), stiffer RBCs tend to get stuck at the apex due to cell collisions and can lead to more intense lingering instead (see “BIF-c” in Fig. 7(*b*)). The distinct effect of RBC deformability on its lingering behavior at the Y-type and T-type bifurcations observed here is in line with a recent report on capillary vascular network [35].

On the other hand, for identical flow conditions, enriched hematocrit (*H*_F_ = 33%) in the mother branch has a weakening effect on RBC lingering at the bifurcation whereas reduced hematocrit (*H*_F_ = 10%) has a strengthening effect, compared to the experimentally measured value *H*_F_ = 26% for the same bifurcation (Fig. 7(*c*)). The probable reason behind the decreased *Pe_λ_* in this case is that crowded cell traffic reduces the possibility of individual RBCs residing on the apex for extended periods of time as they are more likely to be pushed forward by following cells (“herding” effect as reported in [36]). We also considered a different RBC size given the variability of cell morphology *in vivo*, which is modelled through increasing the default cell diameter *D*_RBC_ = 6*μ*m from *D*_RBC_ = 6*μ*m to *D*_RBC_ = 8*μ*m while maintaining the same *H*_F_ (Fig. 7(*c*)). The increased RBC size leads to higher-level of cell-confinement in the capillaries and reduces the fluctuation amplitude of RBC lingering caused by hematocrit alteration.

## Conclusion

Our study demonstrates that lingering (i.e. cells temporarily residing near the bifurcation apex with diminished velocity) is a mechanism which can significantly modify the partitioning of red blood cells (RBCs) through bifurcations. Therefore, it should be taken into account when predicting the hematocrit distribution in microvascular networks at the capillary level.

Existing literature has extensively tested and validated the Zweifach-Fung effect in microvascular bifurcations at the arteriolar level, mostly through microfluidic experiments or numerical simulations. Nonetheless, studies that experimentally quantify the effect of *in vivo* RBC-bifurcation interactions are rare and it remains largely unknown how well the Pries-Secomb model describes the RBC partitioning at the excessively-confined capillary level [35], where the cell-free layer (a key constituent in the empirical model of Pries et al.) becomes negligible and the cells virtually squeeze through vessels in close contact. Previous numerical-experimental works suggested that the branching geometry and cross-sections of the daughter branches can bias the partitioning of capsules at a bifurcation, depending on the size and deformability of the capsule [37]. With much higher confinement reflecting the capillary blood flow, our study represents the first *in vivo* demonstration of the effect of bifurcation geometry and cell properties, and provides a mechanistic justification through the microscopic cell-wall interaction, namely lingering, on the partitioning of RBCs.

Furthermore, we have shown that RBC lingering is inherently associated with certain degree of asymmetry between the daughter branches (Fig. 5), which can either reinforce the Zweifach-Fung effect, or revert it. Indeed, Supp. Fig. S6 illustrates that both positive and negative deviations from the empirical model by Pries et al. can be observed at a high lingering Péclet number. A similar trend is observed for deviations from the linear scaling (which hypothesizes that the fractional erythrocyte flow rate equals the fractional blood flow rate *FQ_e_* = *FQ_b_*, see Supp. Fig. S5). The key finding is that the lingering phenomenon is a prevalent mechanism in the capillary network, subject to fine-tuning of the fractional blood flows and geometrical features of the bifurcation. We demonstrate that, under the assumption of plug flow given highly-confined cell motion, the lingering Péclet number (*Pe_λ_*) correlates with the flow rate ratio *Q_SD_*/*Q_MD_* (*MD* and *SD* refer to the main daughter and secondary daughter branches, respectively). This ratio *Q_SD_*/*Q_MD_* determines the position of the stagnation point, where the local curvature is further associated with the lingering intensity.

Our findings on the modulating effect of RBCs lingering on hematocrit distribution have important implications for remodeling of the microvascular network occurring in development or reintroduced in disease. Recently, it has been uncovered that RBC distribution is strongly associated with the pattern of vascular remodeling, presumably by affecting the wall shear stress difference in neighboring branches through effective blood viscosity [16]. Since lingering can substantially influence the RBC partitioning at capillary bifurcations (in extreme cases revert it), one can expect that lingering may play a key role in shaping the microcirculatory network. Additionally, since the deformability of RBCs affects their characteristic lingering time and, consequently, their partitioning, lingering might be a route through which the microvascular blood flow is impaired in diseases involving hardening of RBCs [23, 24, 25, 26, 27, 28]. Further *in vivo* investigations with rigidified RBCs should be performed in future work to test this hypothesis experimentally.

Finally, this work highlights the need for more models dedicated to RBC studies considering high confinement, as observed in microcirculatory environments entailing the lingering phenomena. Our results highlight the importance of input parameters, such as the flow split in the bifurcation and the curvature of the apex region of the bifurcation.

## Author Contributions

AD, CW, MWL, LK and MDM designed the research. GS, MWL and MDM designed, applied and obtained authorizations for animal experiments. GS and MWL performed the surgery and recorded the movies. YR, AD, AK, MB and TJ designed, implemented and curated the analysis methods for experimental movies. QZ designed the simulation and performed the numerical analysis with input from TK and MOB. YR, AD, QZ and GS wrote the manuscript. All authors discussed the results and critically reviewed the article.

## Acknowledgments

This work was supported by the research unit FOR 2688 Wa1336/12 and LA2682/9-1 of the German Research Foundation, and by the Marie Sklodowska-Curie grant agreement No. 860436—EVIDENCE. A.D. acknowledges funding by the Young Investigator Grant of the Saarland University. Q.Z., T.K. and M.O.B are sponsored by the UKRI Engineering and Physical Sciences Research Council (EPSRC EP/T008806/1). Supercomputing time on the ARCHER2 UK National Supercomputing Service (http://www.archer2.ac.uk) was provided by the ”UK Consortium on Mesoscale Engineering Sciences (UKCOMES)” under EPSRC Grant No. EP/R029598/1, with computational support from the ”Computational Science Centre for Research Communities (CoSeC)” through UKCOMES.

