## supplemental material text and figures for "Red blood cell lingering modulates hematocrit distribution in the microcirculation"

[1]Experimental Physics, Saarland University, 66123 Saarbruecken, Germany
[2]Institute for Clinical and Experimental Surgery, Saarland University, 66421 Homburg, Germany
[3]School of Engineering, Institute for Multiscale Thermofluids, University of Edinburgh, Edinburgh EH9 3FD, United Kingdom
[4]Université Grenoble Alpes, CNRS, LIPhy, F-38000 Grenoble, France
[5]LaMCScI, Faculty of Sciences, Mohammed V University of Rabat, Rabat 1014, Morocco
[6]Centre for Medical Informatics, Usher Institute, University of Edinburgh, Edinburgh EH16 4UX, United Kingdom
[7]The Bayes Centre, The University of Edinburgh, Edinburgh EH8 9BT, United Kingdom
[8]Theoretical Medicine and Biosciences, Saarland University, 66421 Homburg, Germany
[9]Physics and Materials Science Research Unit, University of Luxembourg, L-1511 Luxembourg, Luxembourg
[†]These authors contributed equally to this work.

August 31, 2022

**Abstract**

This document presents supplementary material for the article mentioned in the title.

#### Supplemental Figures and legends of Supplemental Movies

This section regroups caption of the Supplemental Movies and the Supplementary Figures described in the main text.

**Supplemental Movie** M1: (Lingering effect.avi) Recorded movie at a representative bifurcation (Fig.2(a)). The mother vessel (M) and two daughter vessels (MD, SD) are annotated in Fig.2(a). We observed in this video the SD vessel, i.e. with the lowest total blood flow rate, collects more RBCs, in contradiction to the classical partitioning prediction (Zweifach-Fung effect). In addition, RBCs go to the MD vessel almost only if have some interaction with other cells at the apex.

**Supplemental Movie** M2: (SupplementaryMovie_simulation_speedup.mp4) Compilation of animation videos from the numerical simulations performed. Note that the simulation videos included in the movie were sped up for demonstration purpose.

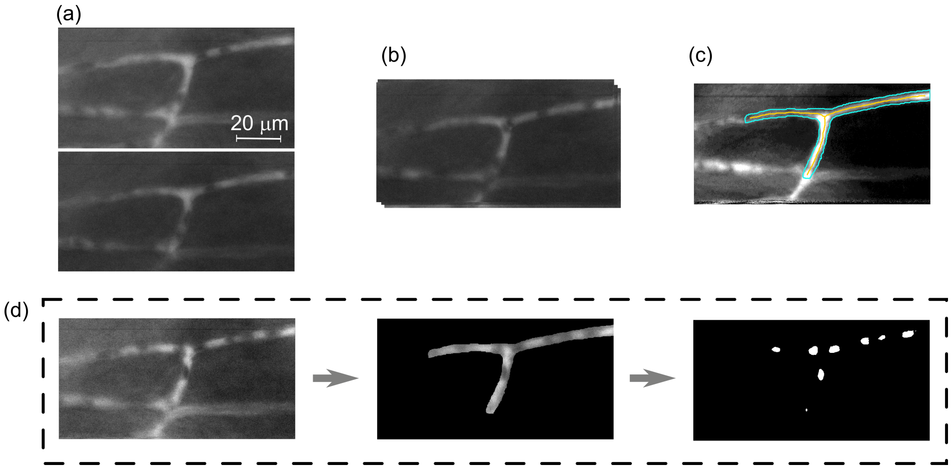


Supplemental Figure **S1**: Schematic representation of the data analysis routine. (*a*) Slight translations of the microscopic field of view, due to the breathing and muscle contraction and expansion of the hamster. (*b*) In order to remove those slight movements in the microscopic field of view, we translated each image by the maximum of its 2d correlation with the first image. A 2D Gaussian filter was applied to remove small defects in each image (to despeckle image series). (*c*) Mask drawn by hand around the vessels. Skeleton pixels of the mask were depicted with the yellow color, using standard Matlab functions (bwmorph). (*d*) The drawn mask was applied to select the Region Of Interest (ROI) of the microscopic field of view and then we binarized the images to detect RBCs.

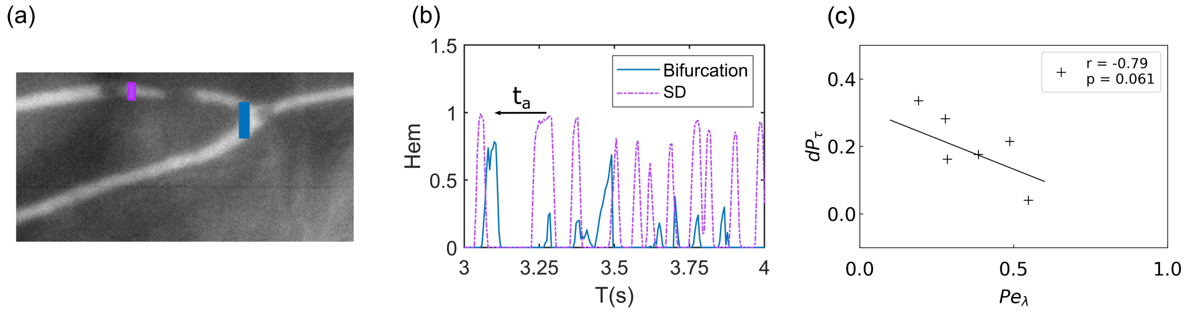


Supplemental Figure **S2**: Extraction of the advection time $t_a$ (*a*) Thin masks used to detect RBCs along time in the bifurcation and one daughter vessel. (*b*) The hematocrit in the masks, extracted from the binarized pictures, is measured along time. The average advection time $t_a$ of the cells in the daughter is obtained by convolution of the two signals. (*c*) The correlation between simulated $dP_\tau$ and $Pe_\lambda$ for "BIF-a", "BIF-b" and "BIF-c" as in Fig. 2 of the main text.

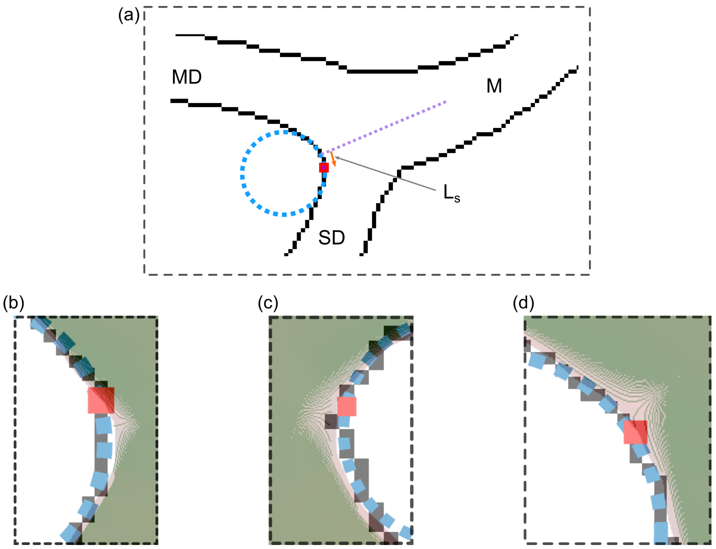


Supplemental Figure **S3**: Experimental extraction of the curvature at the stagnation point, and comparison with numerical simulations. ($a$) Global view of an experiment. The contour of the vessels' mask is in black, while the normal to the bifurcation apex is the (purple) dashed line. The experimentally determined stagnation point is the red square, while its distance at the apex $L_s$ is highlighted with the arrow. The letters $M$, $MD$ and $SD$ refers to the Mother, Main Daughter and Secondary Daughter, respectively. ($b, c, d$) Comparison of the experimental results for the characteristic bifurcations (Fig. 2($a, b, c$)) and the numerical simulations. Obtained stagnation points in experiments and simulations are within one pixel of distance.

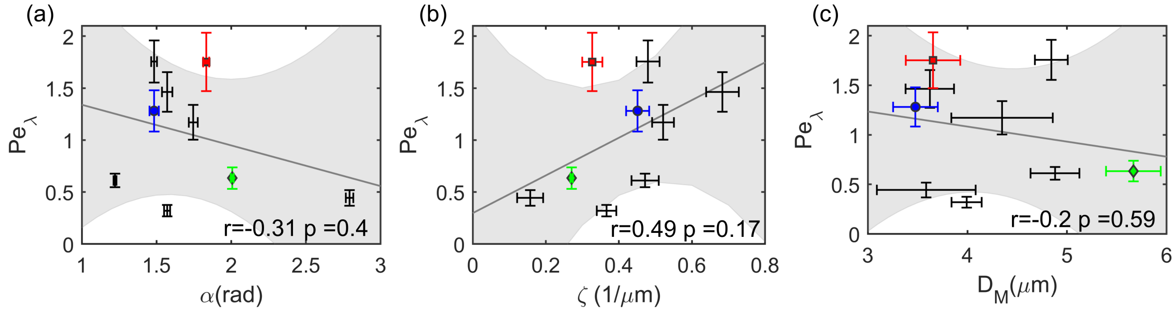


Supplemental Figure **S4**: Bifurcation properties which do not correlate significantly with the lingering Péclet number. (a) opening angle formed by the two daughter vessels. (b) Curvature of the endothelium at the apex of the bifurcation. (c) Diameter of the mother vessel.

### flow divider plane

In our 2D images, the position of stagnation point also coincides with the intersection between the bifurcation wall and the flow divider surface. The flow divider surface is defined as the surface separating the streamlines going into each daughter vessel. As demonstrated by previous studies, in blood vessels the flow divider surface can be efficiently approximated as a plane [1]. The distance between the apex of the bifurcation and this plane, $L_s$, is a function of the flow rate and the flow velocity profile. Since the diameter of the mother branch is smaller than the diameter of the RBC, we assumed here a plug-flow profile, i.e. a velocity distribution mostly flat in the mother vessel. Under these assumptions, the flow divider plane is determined only by the fractional daughter's flow rate [1].

$$\frac{\mathrm{FQ}_{\mathrm{b(MD)}}}{\mathrm{FQ}_{\mathrm{b(SD)}}} = \frac{A_{MD}}{A_{SD}}, \tag{1}$$

Where $A_{MD}$ and $A_{SD}$ are the area to the left and right of the flow divider plane in the mother vessel (see Fig. 6(a), the areas which are depicted with red and blue stripes, respectively). Those can be obtained as half the area of the mother vessel plus (for $A_{MD}$) or minus (for $A_{SD}$) the area between the flow divider plane and the center plane of the vessel. If we assume a cylindrical vessel, one gets

$$\mathrm{A_{MD}} = \frac{\pi R_M^2}{2} + 2\int_0^{L_s} \sqrt{R_M^2 - x^2}\,dx = \frac{\pi R_M^2}{2}\left(1 + \frac{2}{\pi}\arcsin\left(\frac{L_s}{R_M}\right) + \frac{2L_s}{\pi R_M^2}\sqrt{R_M^2 - {L_s}^2}\right), \quad (2)$$

$$\mathrm{A_{SD}} = \frac{\pi R_M^2}{2} - 2\int_0^{L_s} \sqrt{R_M^2 - x^2}\,dx = \frac{\pi R_M^2}{2}\left(1 - \frac{2}{\pi}\arcsin\left(\frac{L_s}{R_M}\right) - \frac{2L_s}{\pi R_M^2}\sqrt{R_M^2 - {L_s}^2}\right), \quad (3)$$

### linear partitioning

The deviation from the empirical model of Pries was calculated and described in the main text of the manuscript (Eq. 2). Similarly, we calculated the deviations from the linear partitioning, $FQ_e = FQ_b$, and results are shown in Fig. **S5**(b) [2, 3, 1]. The Fig. **S5**(b, d) shows the correlation between the lingering Péclet number and various deviations from both behaviors (Eq (2) and linear partitioning). We noted $\Delta_{prediction} = \Delta FQ_e(EX) - \Delta FQ_{e(prediction)}$ the difference of the experimental result with the various predictions (Eq. (2), $ZF$ and $FQ_e = FQ_b$, *linear*). We kept the sign here to see if the lingering effect can enhance ($\Delta_{prediction} > 0$) or decrease ($\Delta_{prediction} < 0$) the heterogeneity partitioning of RBCs, with respect to both Pries model and the linear partitioning. However, both positive and negative deviations are observed at high lingering Péclet number, showing that lingering doesn't systematically favor a specific redistribution of the erythrocytes.

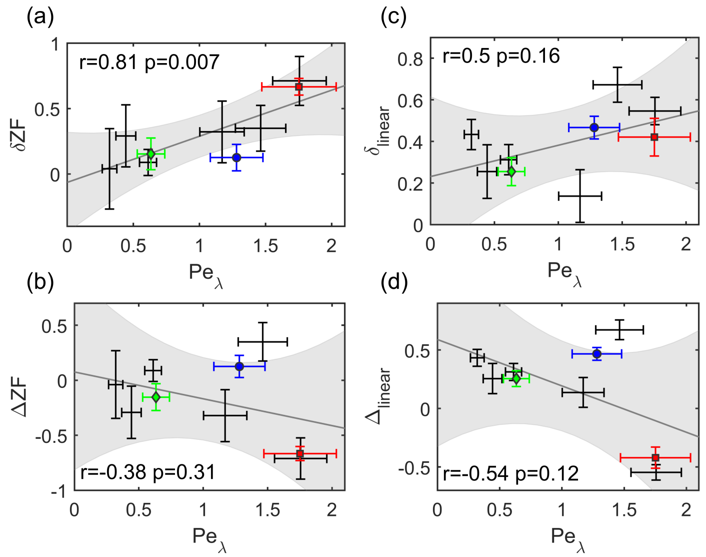


Supplemental Figure **S5**: Deviation of the Pries model and linear partitioning as a function of the lingering Péclet number $Pe_\lambda$. (a), (c) Absolute difference of the experimental result with the prediction of Eq.2 and linear partitioning, respectively. (b), (d) Difference of the experimental result with the prediction of Eq.2 and linear partitioning, respectively (we kept the sign).
